# Extensive N4 Cytosine Methylation is Essential for *Marchantia* Sperm Function

**DOI:** 10.1101/2021.02.12.428880

**Authors:** James Walker, Jingyi Zhang, Yalin Liu, Martin Vickers, Liam Dolan, Keiji Nakajima, Xiaoqi Feng

**Affiliations:** Department of Cell and Developmental Biology, John Innes Centre, Norwich, NR4 7UH, UK; Gregor Mendel Institute of Molecular Plant Biology, Austrian Academy of Sciences, 1030 Vienna, Austria; Nara Institute of Science and Technology, Nara 630-0192, Japan

## Abstract

4-methylcytosine (4mC) is an important DNA modification in prokaryotes, but its relevance, and even presence in eukaryotes have been mysterious. Here we show that spermatogenesis in the liverwort *Marchantia polymorpha* involves two waves of extensive DNA methylation reprogramming. First, 5-methylcytosine (5mC), a well-known eukaryotic DNA modification, expands from transposons to the entire genome. Notably, the second wave installs 4mC throughout genic regions, covering over 50% of CG sites in sperm. 4mC is catalyzed by two novel methyltransferases (MpDN4MT1a and MpDN4MT1b) specifically expressed during late spermiogenesis. Deletion of Mp*DN4MT1s* eliminates 4mC, alters the sperm transcriptome, and produces sperm with swimming defects. Our results reveal extensive 4mC in a eukaryote and define a new family of eukaryotic methyltransferases, thereby expanding the repertoire of functional eukaryotic DNA modifications.

## Main Text

Cytosine methylation plays essential regulatory roles in prokaryotic and eukaryotic genomes (*1, 2*). Both C5 and N4 positions of cytosine can be methylated, forming 5mC and 4mC, respectively (*3*). 4mC is well-documented in bacteria and archaea, but the existence of 4mC in eukaryotic DNA is disputed because the amounts detected are below the technical false positive rates (*4, 5*). Moreover, no eukaryotic N4 DNA methyltransferases have been identified (*6*). 5mC is prevalent across eukaryotes and catalyzed by DNA methyltransferases (DNMT1 and 3) that are highly conserved (*6–8*). Genome distribution of 5mC can be global or mosaic. In vertebrates, 5mC occurs across the whole genome in the CG dinucleotide context, for example covering >80% of CGs in human tissues (*9*). In contrast, flowering plants have mosaic 5mC restricted to transposable elements (TEs) and some genes, occurring in all sequence contexts (CG, CHG and CHH; H = A, C or T) (*10, 11*). 5mC remains largely constant during animal and plant development and exerts homeostatic functions, the disruption of which causes developmental abnormalities and diseases such as cancer (*9, 10, 12, 13*). Notably, 5mC reprogramming occurs and plays essential regulatory roles during spermatogenesis in mammals and flowering plants (*13–16*).

*Marchantia polymorpha* is an early-diverging land plant that shared a last common ancestor with flowering plants approximately 450 million years ago (*17*). The vegetative thallus (a leaf-like tissue) displays similar 5mC patterns to flowering plants, with methylation in the CG, CHG and CHH contexts associated with TEs (*18, 19*) (16% of the genome is methylated; Fig. 1A). However, unlike flowering plants, in which some genes are CG methylated, *Marchantia* vegetative tissues have no methylation in genes, similar to other early-diverging land plant species, *Physcomitrella patens* and *Selaginella moellendorffii* (*7, 18, 20*). In contrast to the mosaic methylation in vegetative tissues, global cytosine methylation (covering 97% of the genome; Fig. 1A) has been reported in *Marchantia* sperm in all sequence contexts (*19*). The switch from mosaic methylation patterns in vegetative thallus to the global pattern in sperm suggests that genome-scale DNA methylation reprogramming occurs during male reproductive development.

**Fig. 1.**
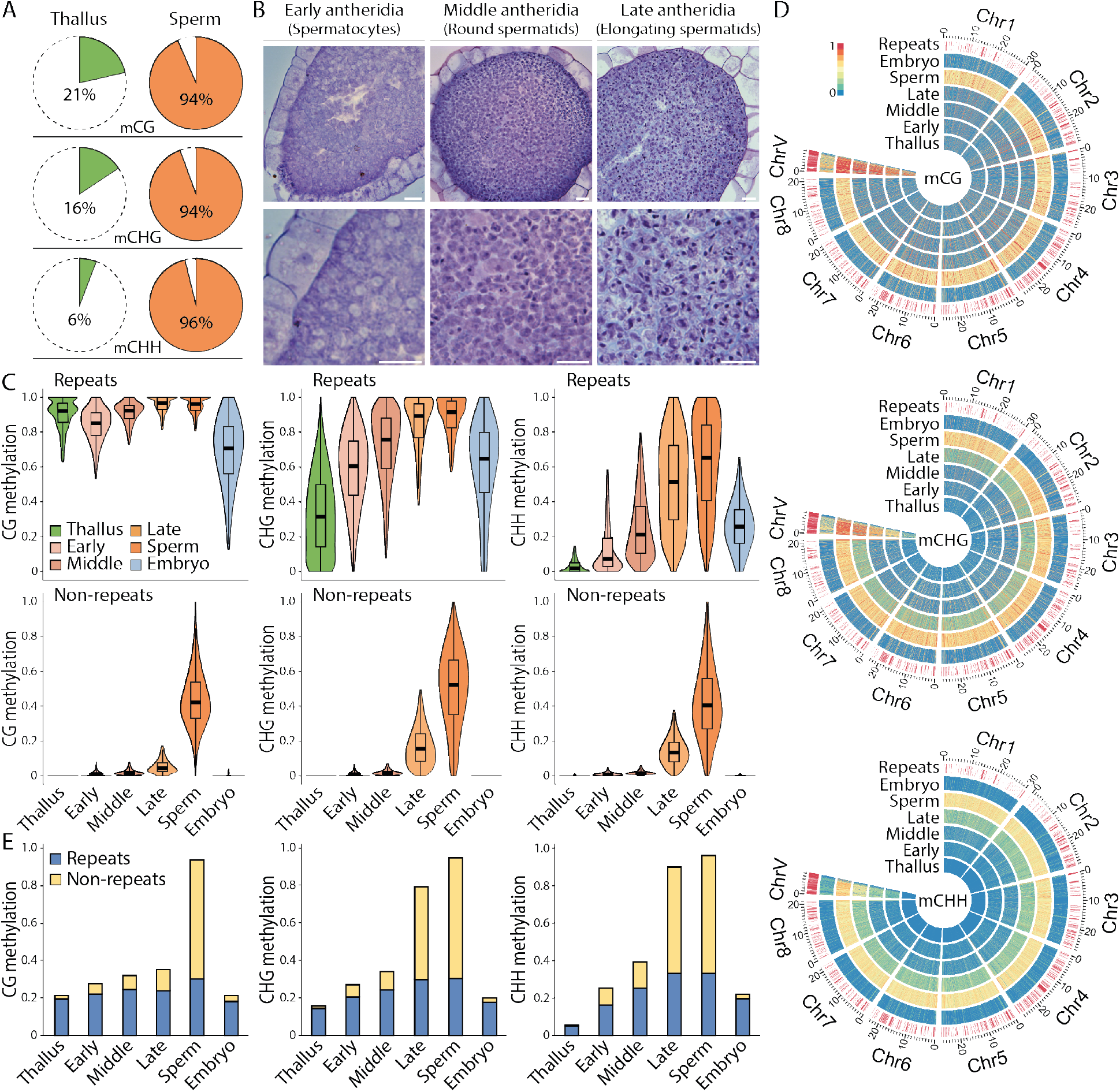
Two waves of DNA methylation reprogramming occur during *Marchantia* sperm development. (**A**) Pie charts illustrating percentage of 100-bp windows across the *Marchantia* genome with > 0.2 CG methylation, > 0.1 CHG methylation, or > 0.05 CHH methylation in thallus and sperm. (**B**) Transverse sections of developing antheridia stained with toluidine blue. Scale bars, 20 μm. (**C**) Violin plots showing methylation for 100-bp windows associated with repeats or non-repeats for thallus, antheridia (of early, middle and late stage), sperm and embryo. (**D**) Heat maps displaying methylation for 10-kb windows across the *Marchantia* Tak-1 chromosomes (Chr1-8, autosomes; ChrV, male sex chromosome) in the tissues shown in (C). Red bars indicate 10-kb windows covered by > 95% TEs. (**E**) Bar graphs depicting percentages of 100-bp genomic windows with evident methylation (as in (A)) in various tissues. Blue and yellow show windows associated with repeats and non-repeats, respectively. Early, middle and late, represent early, middle and late antheridia, respectively (C to E).

To identify the mechanism of cytosine methylation reprogramming in *Marchantia* sperm, we isolated mature sperm as well as antheridia (male organs analogous to the testes). Antheridia were staged by microscopy and separated into early (containing spermatocytes), middle (early spermiogenesis, containing mostly round spermatids) and late (late spermiogenesis when DNA compaction and sperm tail formation occur, containing mostly elongating spermatids) stages (Fig. 1B). The developmental staging was confirmed by examining expression of genes with known transcription patterns (*21*) (Fig. S1). To inspect methylation following fertilization, we also isolated embryos.

Bisulfite sequencing (BS-seq; Table S1) of developing antheridia and mature sperm showed that cytosine methylation at TEs increases considerably throughout spermatogenesis in all sequence contexts, from 38% average fractional methylation in early antheridia to 72% in mature sperm (27% in thallus; Fig. 1C and Fig. S2). This is most evident in the male sex chromosome (Chromosome V) because of the high density of TEs on this chromosome (*22*) (Fig. 1D). We also observed two temporal waves of methylation expansion into the non-repetitive parts of the genome. The first wave involves non-CG (CHG and CHH) methylation, which expands throughout spermiogenesis from covering 35% of the genome in round spermatids (middle antheridia stage) to 80% in elongating spermatids (late antheridia stage) and ultimately to 96% in mature sperm (Fig. 1, C to E, and Fig. S2). The second wave manifests in the expansion of CG methylation primarily at the last stage of spermiogenesis (late antheridia onwards; Fig. 1, C to E, and Fig. S2). This event culminates in methylation of 94% of the genome in mature sperm, compared to 35% in late antheridia and 21% in thallus (Fig. 1E). Methylation expansion at this scale has only been previously reported in the mammalian germline, where DNA methylation is re-established genome-wide following global demethylation in primordial germ cells (*15, 16*). Methylation patterns in the embryo are similar to those of thallus, with an absence of both CG and non-CG methylation at genic regions (Fig. 1, C to E, and Fig. S2), indicating that sperm hypermethylation is fully reversed in the zygote or during embryo development.

An examination of transcriptomic dynamics during spermatogenesis revealed an increase in the expression of DNA methyltransferase Mp*DNMT3b* and chromomethylase Mp*CMTa* (Fig. 2A and Table S2), orthologs of which are involved in CHH and CHG methylation of TEs in *P. patens* (*23, 24*). The temporal expression patterns of these enzymes correlate with the timing of TE methylation reinforcement and non-CG methylation expansion (Fig. 1, C to E, Fig. 2A, and Fig. S2), suggesting the likely involvement of Mp*DNMT3b* and Mp*CMTa* in these events. To test this hypothesis we generated Mp*dnmt3b* and Mp*cmta* knockdown mutants and measured methylation in their sperm. CHG and CHH methylation throughout the genome is greatly reduced in the sperm of both Mp*dnmt3b* and Mp*cmta* knockdown mutants compared to wild type, whereas CG methylation remains largely unchanged (Fig. 2, B to D, and Fig. S3). Therefore, MpDNMT3b and MpCMTa mediate non-CG methylation reprogramming in sperm.

**Fig. 2.**
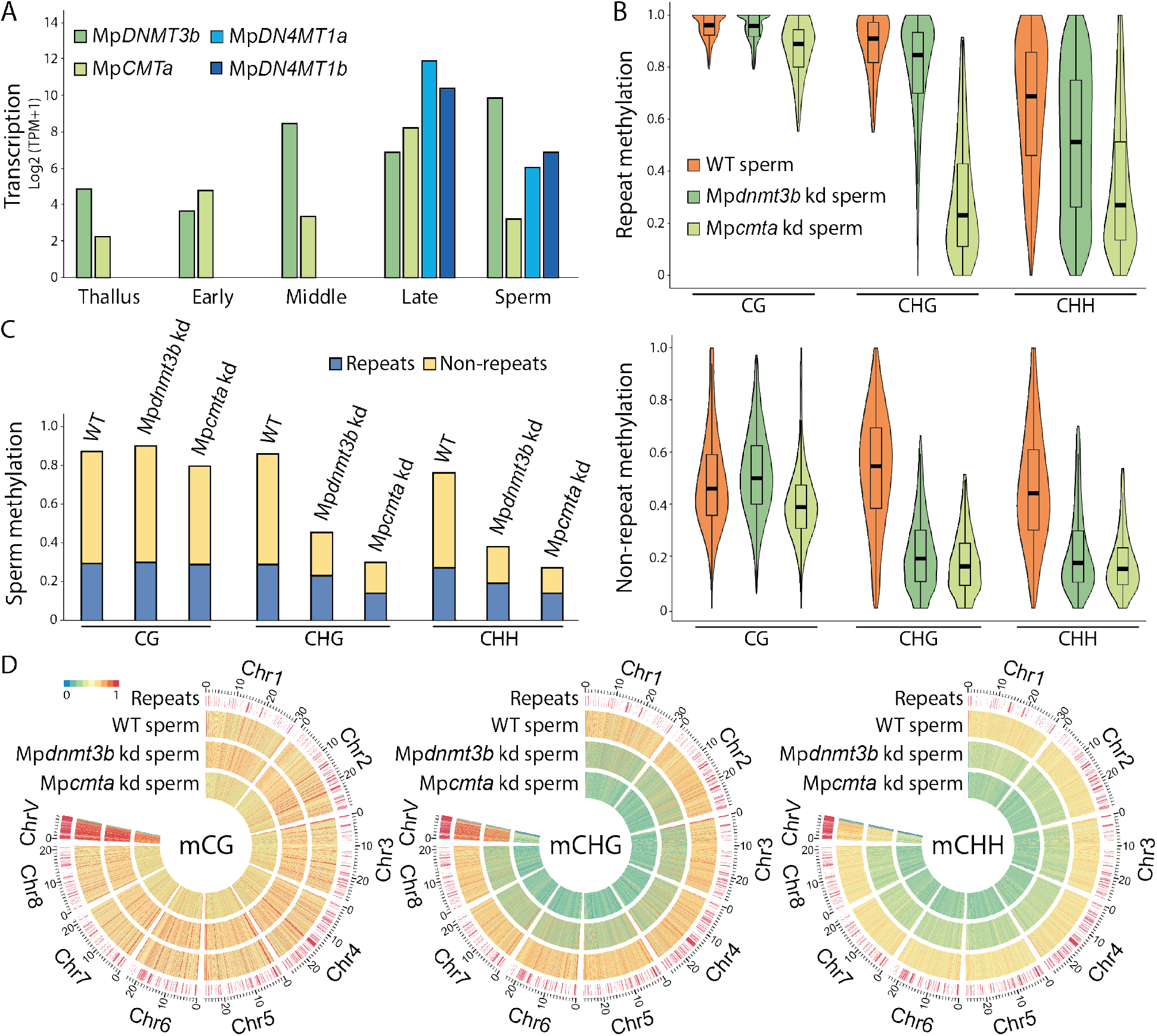
Non-CG methylation reprogramming in sperm requires Mp *DNMT3b* and Mp *CMTa*. (**A**) Transcript levels of Mp*DNMT3b*, Mp*CMTa*, Mp*DN4MT1a* and Mp*DN4MT1b* in the thallus, antheridia of early, middle and late stages, and sperm. TPM, transcripts per million. (**B**) Violin plots showing CG, CHG or CHH methylation at repeats or non-repeats, as in Fig. 1C, in the sperm of the wild-type (WT), the Mp*dnmt3b* knockdown (kd) mutant and the Mp*cmta* kd mutant. (**C**) Bar graphs illustrating the percentage of 100-bp genomic windows with > 0.3 methylation in CG, CHG or CHH context in WT, Mp*dnmt3b* kd, and Mp*cmta* kd sperm. Blue and yellow show windows associated with repeats and non-repeats, respectively. (**D**) Heat maps as in Fig. 1D showing methylation for WT, Mp*dnmt3b* kd, and Mp*cmta* kd sperm.

The timing of CG methylation expansion suggests that it is caused by enzyme(s) that are expressed at the last stage of sperm maturation. Since none of the known *Marchantia* methyltransferases – including Mp*MET*, the homolog of the CG methyltransferase *DNMT1* (*25*) – has such an expression pattern (Table S2), we examined all genes expressed during late spermiogenesis. Among the most highly transcribed were two tandemly duplicated genes (Mp6g18330 and Mp6g18340; Fig. 2A and Table S2) that cluster with prokaryotic N4 cytosine methyltransferases in a phylogenetic tree (Fig. S4A) and contain TSPPY sequences characteristic of the N4 cytosine methyltransferase catalytic domain (*26, 27*) (Fig. S4B). The arrangement of motifs in these proteins also reflects prokaryotic 4mC methyltransferases rather than 5mC methyltransferases (*26, 28*) (Fig. S4B). Transcripts of both genes are absent in the thallus and published datasets from other tissues (*17*) and only detected in late antheridia containing elongating spermatids and in the mature sperm (Fig. 2A and Table S2). Mp6g18330- and Mp6g18340-citrine fusion proteins expressed under native promoters confirm the specific expression in elongating spermatids (Fig. S5). We thus hypothesized that these proteins catalyze the formation of 4mC that contributes to the global CG methylation expansion observed during late spermiogenesis. These genes will hereafter be referred to as *Marchantia polymorpha DNA N4 CYTOSINE METHYLTRANSFERASE1a* (Mp*DN4MT1a*) and Mp*DN4MT1b*.

To test our hypothesis, we quantified modified nucleotides present in the DNA of thallus and mature sperm by liquid chromatography–mass spectrometry (LC-MS). The 5mC content of thallus DNA (7.6%) is consistent with our BS-seq results, and there is no detectable 4mC (Fig. 3, A and B). Mature sperm has much higher levels of 5mC (31.0%), in agreement with the hypermethylation observed by BS-seq (Fig. 3B). However, this is substantially less than the 52.7% cytosine methylation measured by BS-seq (Fig. 3B), a technique that detects 4mC as well as 5mC (*3, 29*). Consistent with our hypothesis on the existence of 4mC in *Marchantia* sperm, LC-MS revealed that 4mC comprises a striking 15.9% of the cytosines in sperm (Fig. 3, A and B), far exceeding 4mC levels previously detected in prokaryotes (typically <1%) (*4*). To test the link between MpDN4MT1s and 4mC, we generated a Mp*dn4mt1a* and Mp*dn4mt1b* double knockout mutant (simplified as Mp*dn4mt1*) via CRISPR/Cas9-mediated deletion (Fig. S6) and performed LC-MS on isolated sperm. As expected, 4mC is undetectable in the Mp*dn4mt1* mutant (Fig. 3, A and B), demonstrating that MpDN4MT1s are indeed N4 cytosine methyltransferases that catalyze extensive 4mC in *Marchantia* sperm.

**Fig. 3.**
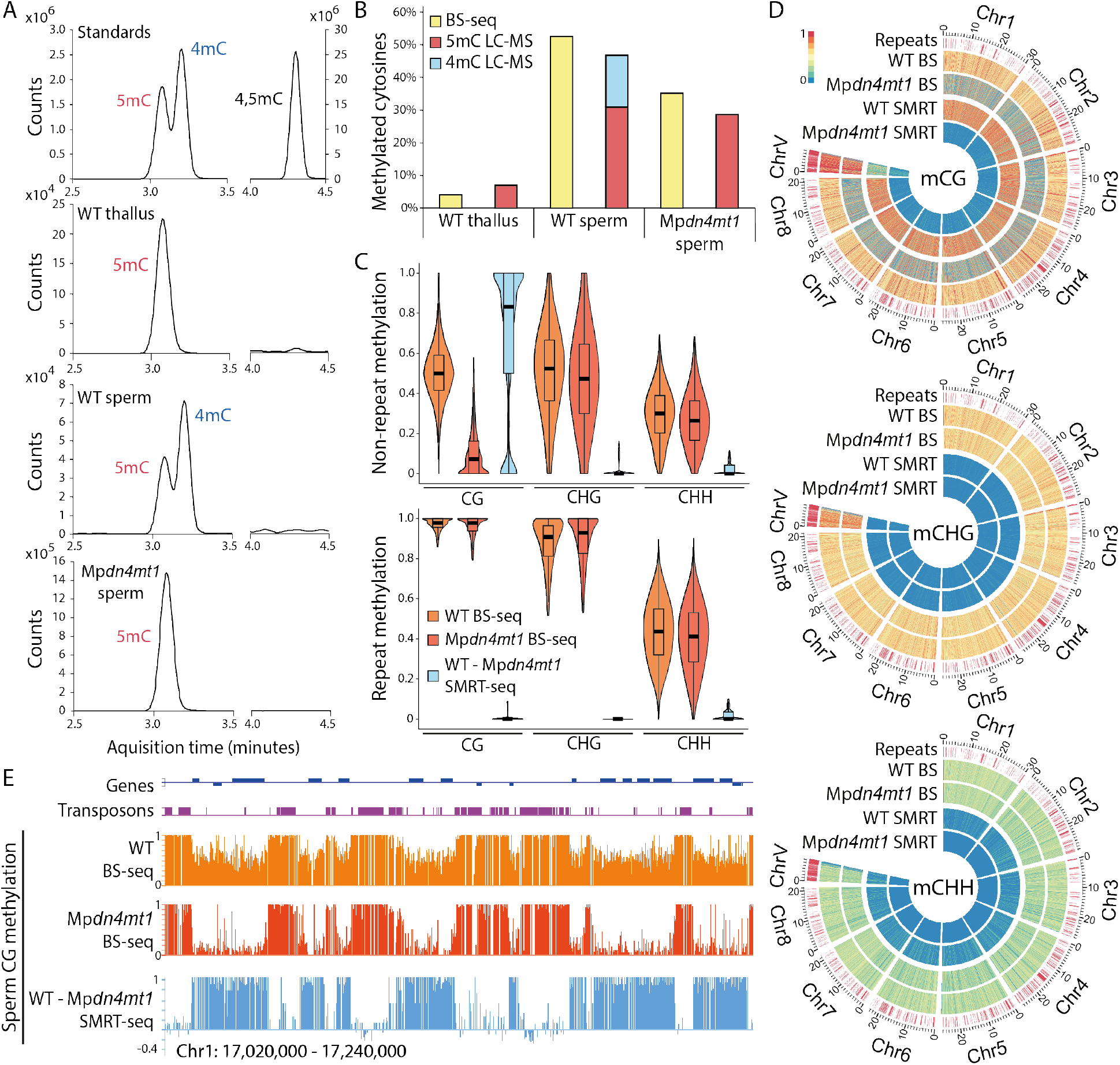
MpDN4MT1s catalyze extensive 4mC across genic regions in sperm. (**A**) LC-MS peaks of methylated deoxycytidine (5mC, 4mC and 4,5mC) standards and DNA isolated from WT thallus, WT sperm, and Mp*dn4mt1* mutant sperm. (**B**) Percentages of methylated cytosines in WT thallus, WT sperm, and Mp*dn4mt1* sperm detected by BS-seq or LC-MS. (**C**) Violin plots as in Fig. 1C showing methylation of 100-bp genomic windows detected by BS-seq in WT and Mp*dn4mt1* sperm, and 4mC detected by SMRT-seq in WT sperm minus Mp*dn4mt1* sperm, associated with either repeats or non-repeats. (**D**) Heat maps as in Fig. 1D, depicting methylation in WT and Mp*dn4mt1* sperm detected by either BS-seq or SMRT-seq. (**E**) Snapshots of CG methylation detected by BS-seq of WT and Mp*dn4mt1* sperm, as well as SMRT-seq (showing WT minus Mp*dn4mt1* sperm methylation).

BS-seq utilizes the resistance conveyed by the methyl group to bisulfite-mediated deamination of cytosine to uracil (*30*). Unlike 5mC, which is almost completely resistant to bisulfite conversion, 4mC is only partially resistant with roughly 43-55% of 4mC remaining unmodified in a typical treatment (*29, 31*). To investigate the context and distribution of N4 cytosine methylation within the sperm genome, we performed BS-seq with one round of bisulfite treatment, and compared to data obtained from two treatments, as we expect 4mC to be progressively deaminated while 5mC remains unmodified. This comparison revealed a sensitivity of CG methylation to the second treatment in non-repetitive genomic regions (16% methylation loss, similar to 14% methylation reduction of a 4mC spike-in control), but minimal sensitivity of CG methylation in repeats (1.8%) and similarly negligible sensitivity in non-CG contexts throughout the genome (1.7%; Fig. S7). Consistently, sequencing of Mp*dn4mt1* knockout sperm with one bisulfite treatment shows a near-complete loss of CG methylation at non-repetitive genic regions, whereas non-CG methylation is largely unaltered (Fig. 3, C to E, and Fig. S8). The specific sensitivity of CG methylation in genic regions to bisulfite conversion and its dependence on Mp*DN4MT1*s indicate that genic CG methylation in sperm is catalyzed by MpDN4MT1s.

To further characterize the distribution of 4mC in the sperm genome, we performed PacBio Single Molecule, Real-Time sequencing (SMRT-seq; Table S3), which can distinguish between 4mC and 5mC (*32, 33*). 18% of cytosines were identified as N4 methylated in wild-type, in contrast to 4% in the Mp*dn4mt1* mutant (Table S4). This is comparable to our LC-MS results (Fig. 3B) and consistent with complete dependence of 4mC on Mp*DN4MT1a* and Mp*DN4MT1b*, given the 0.5-7% false discovery rates reported for 4mC detection by SMRT-seq (*5*). SMRT data also confirms that MpDN4MT1s preferentially target CG dinucleotides, with 63% of total CG sites being 4mC methylated (80% outside of TEs), compared to 8% of CHG sites and 8% of CHH sites (Fig. 3, C and D, Fig. S8, and Table S4). This preference is lost in the Mp*dn4mt1* mutant, with 6% 4mC detected at CG sites, 5% at CHG sites, and 4% at CHH sites (Table S4). The apparent false discovery rate of about 4% indicates that N4 methylation of non-CG cytosines is rare, whereas methylation of CG sites in genes and other non-repetitive sequences surpasses 75% (Table S4).

A notable feature of our SMRT-seq data is the infrequent N4 methylation of CG sites in TEs (16%; Fig. 3, D and E, Fig. S8 and Table S4). TEs are extensively C5 CG methylated (Fig. 3C), and our result could be explained by the failure of SMRT-seq to detect N4 methylation at cytosines methylated at the 4^th^ and 5^th^ positions. Alternatively, existing 5mC at CG sites could prevent N4 methylation by MpDN4MT1s during the last stage of sperm maturation. To distinguish these possibilities, we explored the existence of cytosines that are both C5 and N4 methylated (4,5mC) in *Marchantia* sperm by LC-MS, utilizing a synthesized 4,5mC standard (Fig. 3A). This experiment failed to detect any 4,5mC (Fig. 3A), indicating that the MpDN4MT1s do not methylate 5mC to form 4,5mC. Thus, genic regions in *Marchantia* sperm are distinguished by N4 methylation at CG sites, whereas TEs are marked by C5 CG methylation.

The specific expression of MpDN4MT1s during late spermiogenesis suggests that 4mC is important for sperm function. Consistently, global alteration of gene expression is observed in Mp*dn4mt1* mutant sperm (Fig. 4A). In comparison to wild-type, 33% of the transcripts are significantly mis-regulated in Mp*dn4mt1* mutant sperm (Fig. 4A and Fig. S9A), indicating that 4mC regulates gene expression. To evaluate the functional significance of 4mC, we examined the fertility and swimming behavior of wild-type and Mp*dn4mt1* mutant sperm. Crossing female wild-type *Marchantia* with Mp*dn4mt1* mutant sperm shows that mutant sperm can fuse with the egg to produce viable sporophytes (the diploid generation; Fig. S9B). However, the swimming of Mp*dn4mt1* mutant sperm is significantly less directional and slower compared to wild-type sperm (Fig. 4, B and C, and Movies S1 to S4), demonstrating that 4mC is important for sperm function.

**Fig. 4.**
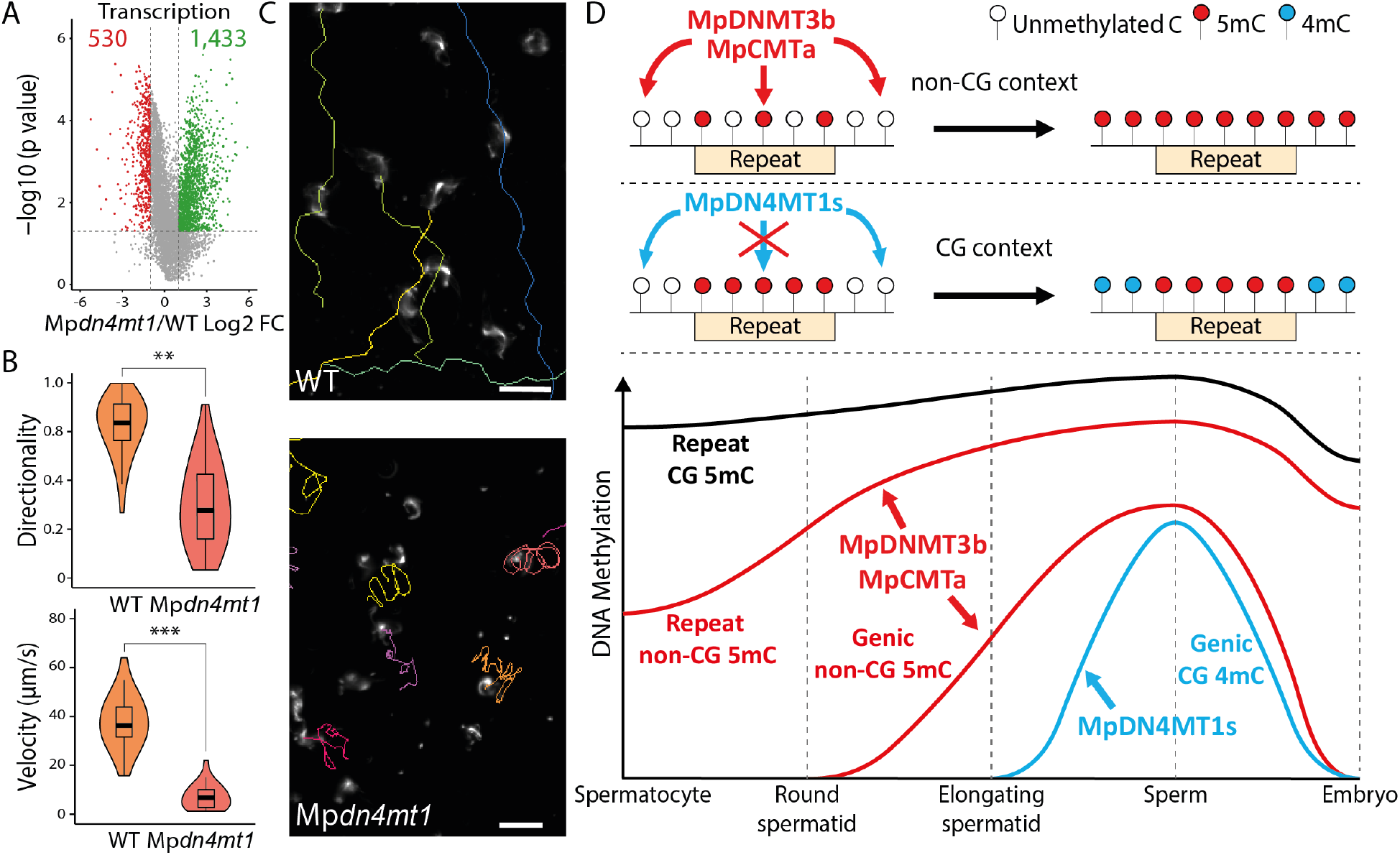
MpDN4MT1-catalyzed 4mC is essential for sperm function. (**A**) Volcano plot illustrating transcripts (TPM > 10 in either genotype) that are upregulated (green) or downregulated (red) in M*pdn4mt1* sperm compared to WT. Vertical dashed lines mark two-fold changes (FC) and the horizontal dashed line shows *P* = 0.05 (likelihood-ratio test). (**B**) Directionality and velocity of motile sperm. *n* = 30 (WT) and 28 (Mp*dn4mt1*). ** *P* < 1.2e^−10^, *** *P* < 4.2e^−15^, Kolmogorov-Smirnov test. (**C**) Snapshots of Movies S1 and S2. Swimming paths taken by example WT or Mp*dn4mt1* sperm are indicated by colored lines. Scale bars, 20 μm. (**D**) Temporal patterns and mechanisms of methylation reprogramming during *Marchantia* sperm development. First, 5mC mediated by MpDNMT3b and MpCMTa in non-CG contexts is reinforced over repeats and expands into genic regions during spermatogenesis. Subsequently during late spermiogenesis, MpDN4MT1s establish 4mC in the CG context across the genome, except in TEs covered by 5mC.

Although 4mC has been extensively characterized in prokaryotes, whether 4mC has any function – or even exists – in eukaryotes is unknown (*4, 5*). Here we show that late spermiogenesis in *Marchantia* involves extensive N4 methylation of genic regions by novel methyltransferases (MpDN4MT1a and MpDN4MT1b) that have a strong *in vivo* preference for CG dinucleotides (Fig. 4D). The specific expression of MpDN4MT1s during the last stage of sperm maturation and their requirement for normal sperm swimming behavior demonstrate the biological importance of 4mC. Overall, our results elucidate the intricate reprogramming of DNA methylation during *Marchantia* spermatogenesis (Fig. 4D) and establish 4mC as a functional DNA modification in eukaryotes.

## Acknowledgments

We thank the John Innes Centre Small Molecule Mass Spectrometry (Lionel Hill) and Chemistry (Martin Rejzek) platforms for assistance with LC-MS, and the Bioimaging Facility (Eva Wegel and Sergio Lopez) for assistance with microscopy. Finally, we thank the Norwich BioScience Institute Partnership Computing infrastructure for Science Group for High Performance Computing resources.

## Author contributions

J.W., J.Z., and X.F. designed the study, J.W., J.Z., and Y.L. performed the experiments, J.W., J.Z., and M.V. analyzed the data, J.W. and X.F. wrote the manuscript.

---

**Fig. S1.**
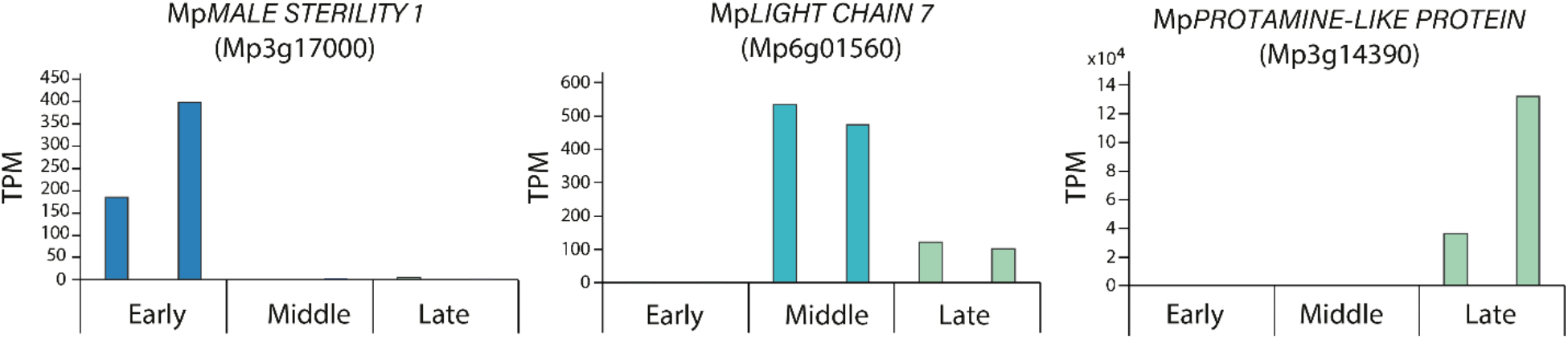
Gene expression during *Marchantia* spermatogenesis. Transcript abundances detected in early, middle or late stage antheridia (each with two biological replicates) for three genes previously known to be developmentally regulated as shown by RNA *in situ* hybridization (*21*). TPM, transcripts per million.

**Fig. S2.**
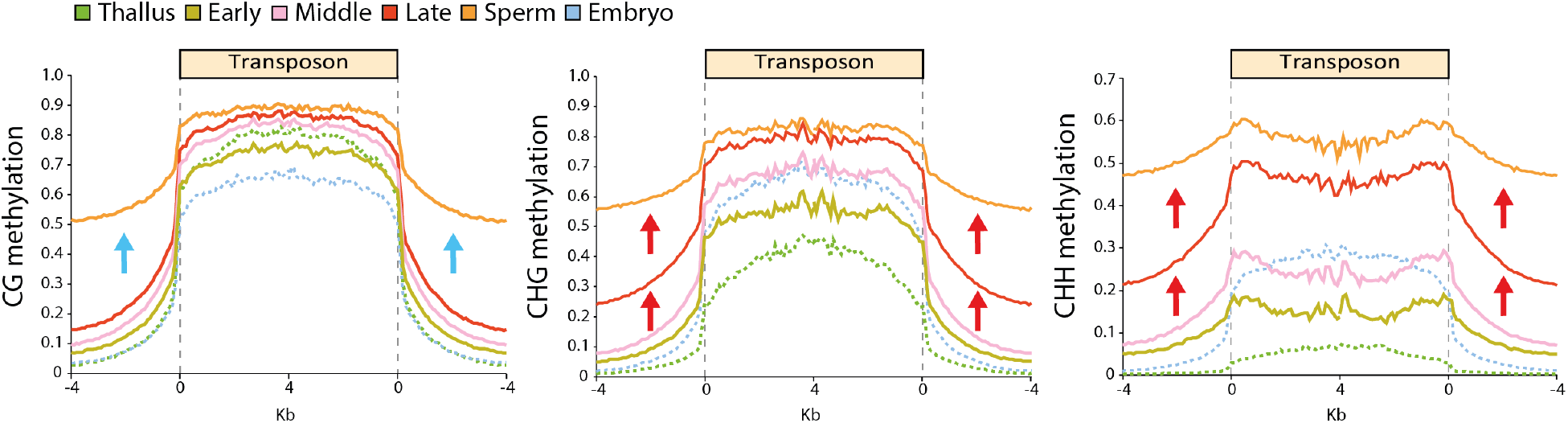
*Marchantia* spermatogenesis features reinforcement of TE methylation and two waves of methylation expansion into non-repetitive regions. DNA methylation over TEs in the antheridia (of early, middle or late stages) and sperm (illustrated by solid lines), in comparison to the thallus and embryo (dashed green and blue lines, respectively). TEs were aligned at the 5’ and 3’ ends (dashed grey lines) and average methylation levels for each 100-bp interval are plotted. Blue arrows indicate expansion of CG methylation to non-repetitive regions at the last stage of sperm maturation (after late antheridia, the red line). Red arrows illustrate expansion of non-CG methylation into non-repetitive regions during spermiogenesis, starting from middle antheridia (pink line).

**Fig. S3.**
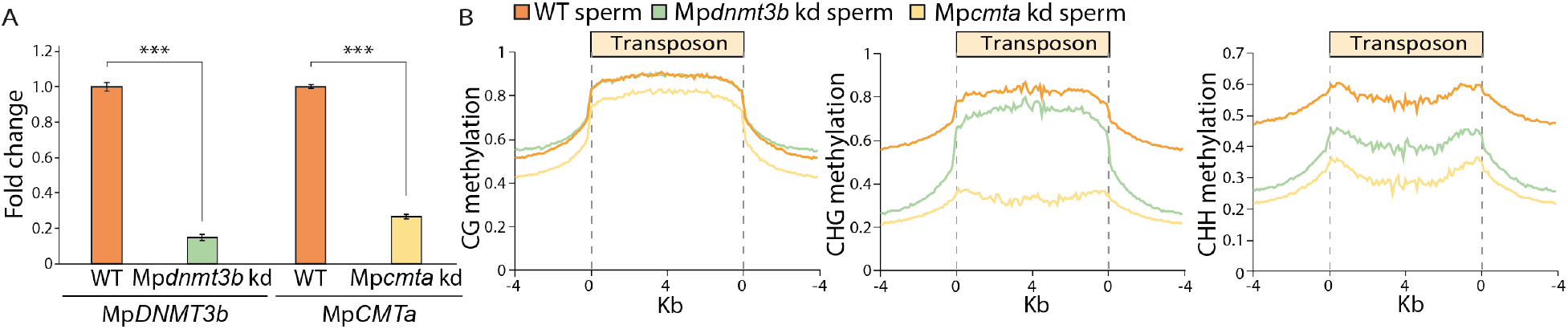
Validation of Mp*dnmt3b* and Mp*cmta* knockdown mutants and their sperm methylation. (**A**) Quantitative RT–PCR showing the expression of Mp*DNMT3b* and Mp*CMTa* in Mp*dnmt3b* kd or Mp*cmta* kd mutant antheridiophores. *** *P* < 0.001, two-tailed *t*-test; *n* = 3. (**B**) Methylation over TEs in WT, Mp*dnmt3b* kd, and Mp*cmta* kd mutant sperm, as in Fig. S2.

**Fig. S4.**
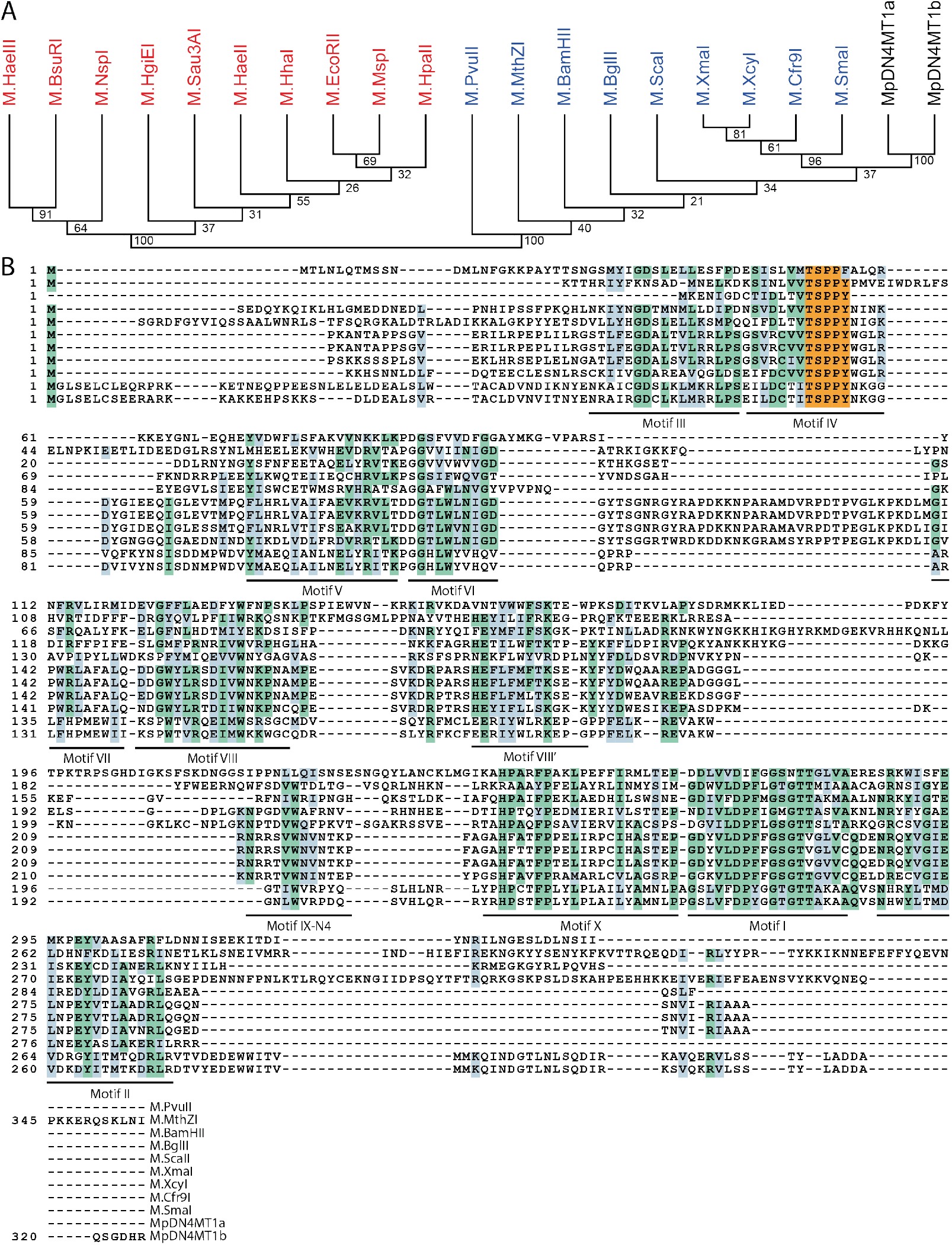
MpDN4MT1a and MpDN4MT1b share homology with prokaryotic 4mC DNA methyltransferases. (**A**) Cladogram of orthologous protein sequences representing prokaryotic 5mC (red) and 4mC (blue) methyltransferases (*26, 28*) with the two MpDN4MT1s in black. (**B**) Protein sequence alignment of MpDN4MT1a and b with the 4mC methyltransferases used in (A). Amino acids with shared identity between sequences are shaded green while similar amino acids are shaded blue. The TSPPY motif characteristic of 4mC methyltransferases is highlighted in orange.

**Fig. S5.**
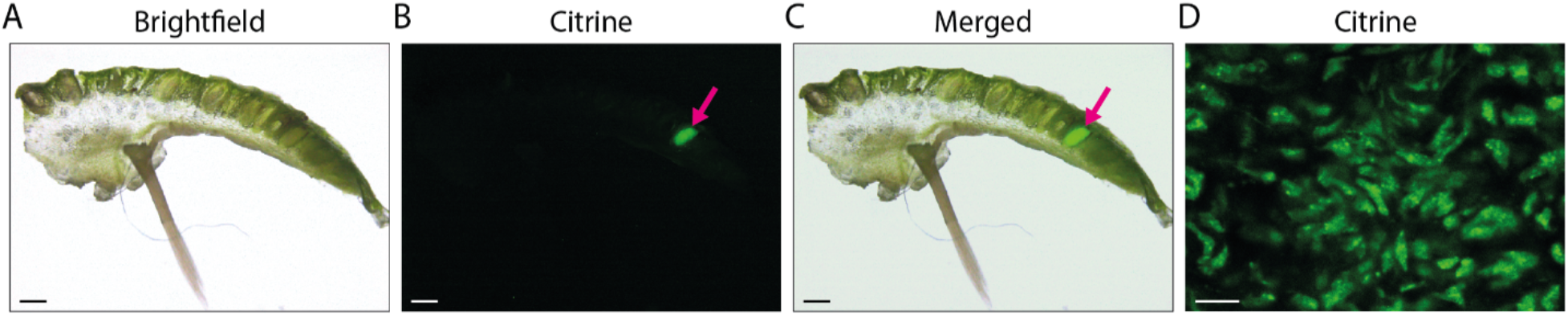
MpDN4MT1a and MpDN4MT1b are specifically expressed in developing sperm. (**A** to **C**) Longitudinal section of Mp*DN4MT1a-Citrine* late-stage antheridiophore using fluorescence microscopy. An antheridium with positive signal is indicated by a magenta arrow. (**D**) Zoomed-in view of elongating spermatids where MpDN4MT1b-Citrine signals are specifically observed via confocal imaging. Scale bars, 500 μm (A to C); 5 μm (D).

**Fig. S6.**
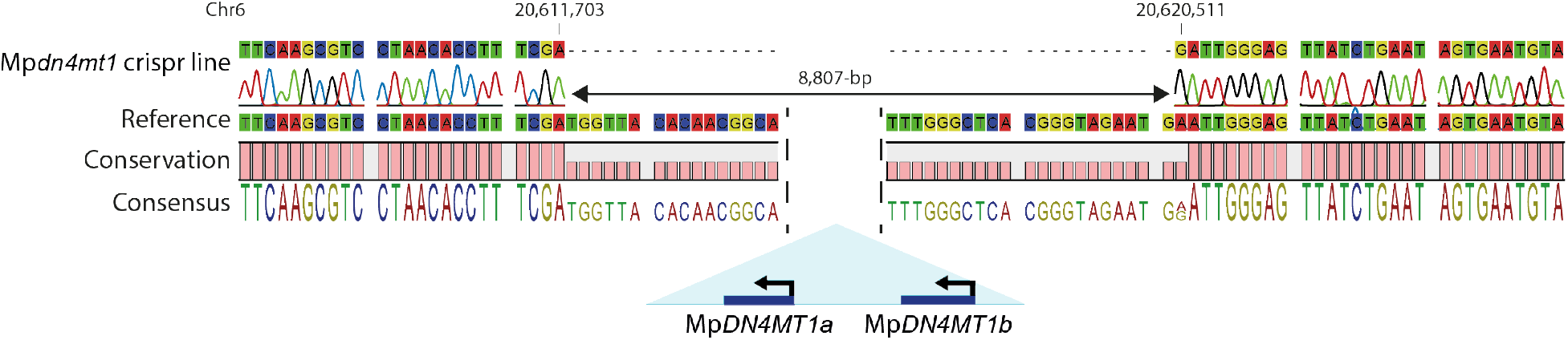
CRISPR/Cas9 deletion of Mp*DN4MT1a* and Mp*DN4MT1b*. DNA sequence alignment illustrating the 5’ and 3’ ends for the CRISPR/Cas9 deletion of Mp*DN4MT1a* and Mp*DN4MT1b* in the Mp*dn4mt1* mutant.

**Fig. S7.**
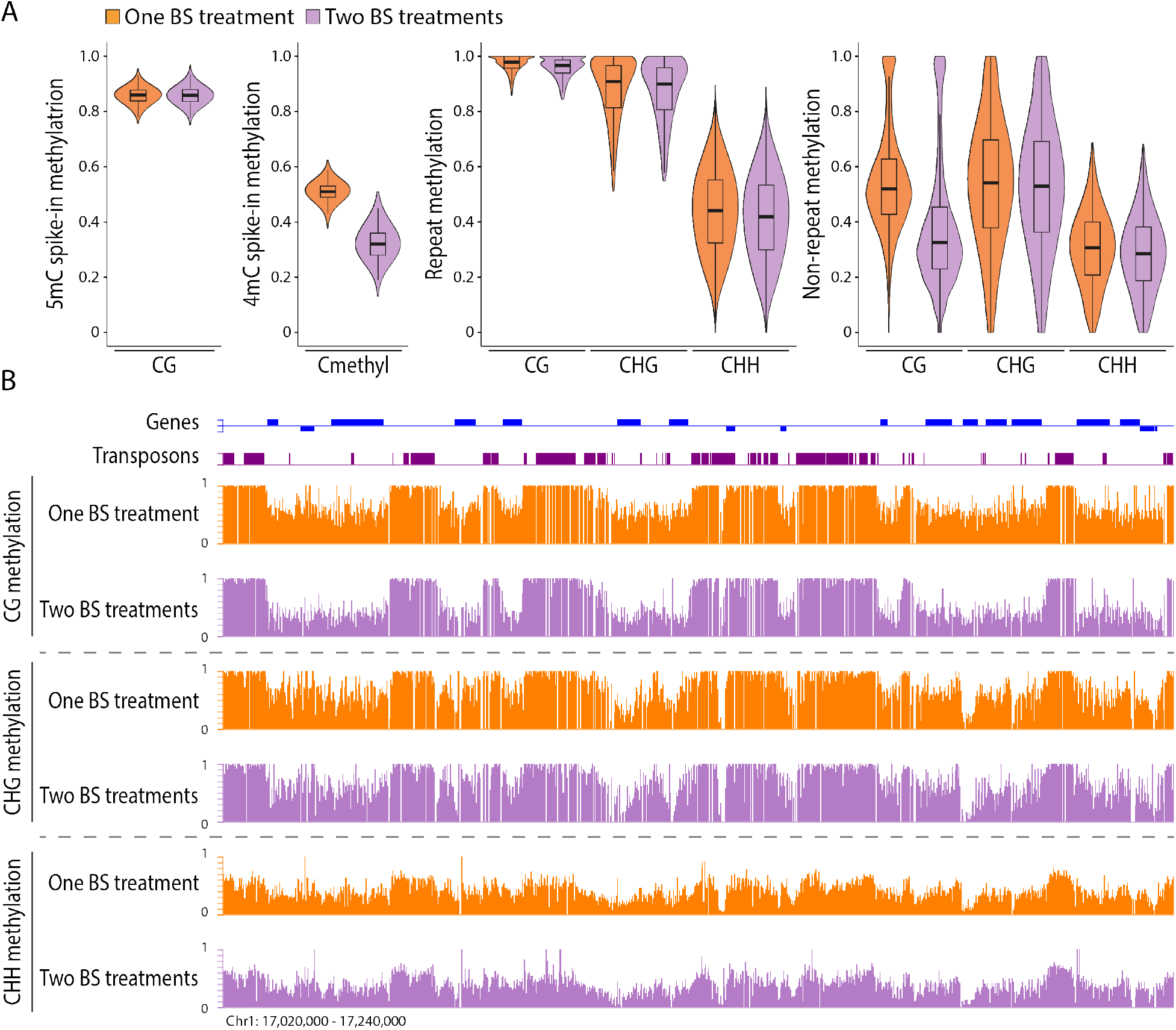
CG methylation at non-repetitive regions is sensitive to bisulfite treatment. (**A**) Violin plots showing DNA methylation detected from BS-seq with one or two rounds of BS treatment of *Marchantia* sperm, with a 5mC spike-in control (CG methylated Lambda genomic DNA) and a 4mC spike-in control (pUC19 PCR product synthesized with 4mdCTP). All cytosine contexts (Cmethyl) are examined for the 4mC spike-in control. Methylation in *Marchantia* sperm is separated into different sequence contexts and genomic regions (repeats and non-repeats). (**B**) Snapshots of methylation levels detected by one or two BS treatments in different cytosine contexts over repeats and genic regions.

**Fig. S8.**
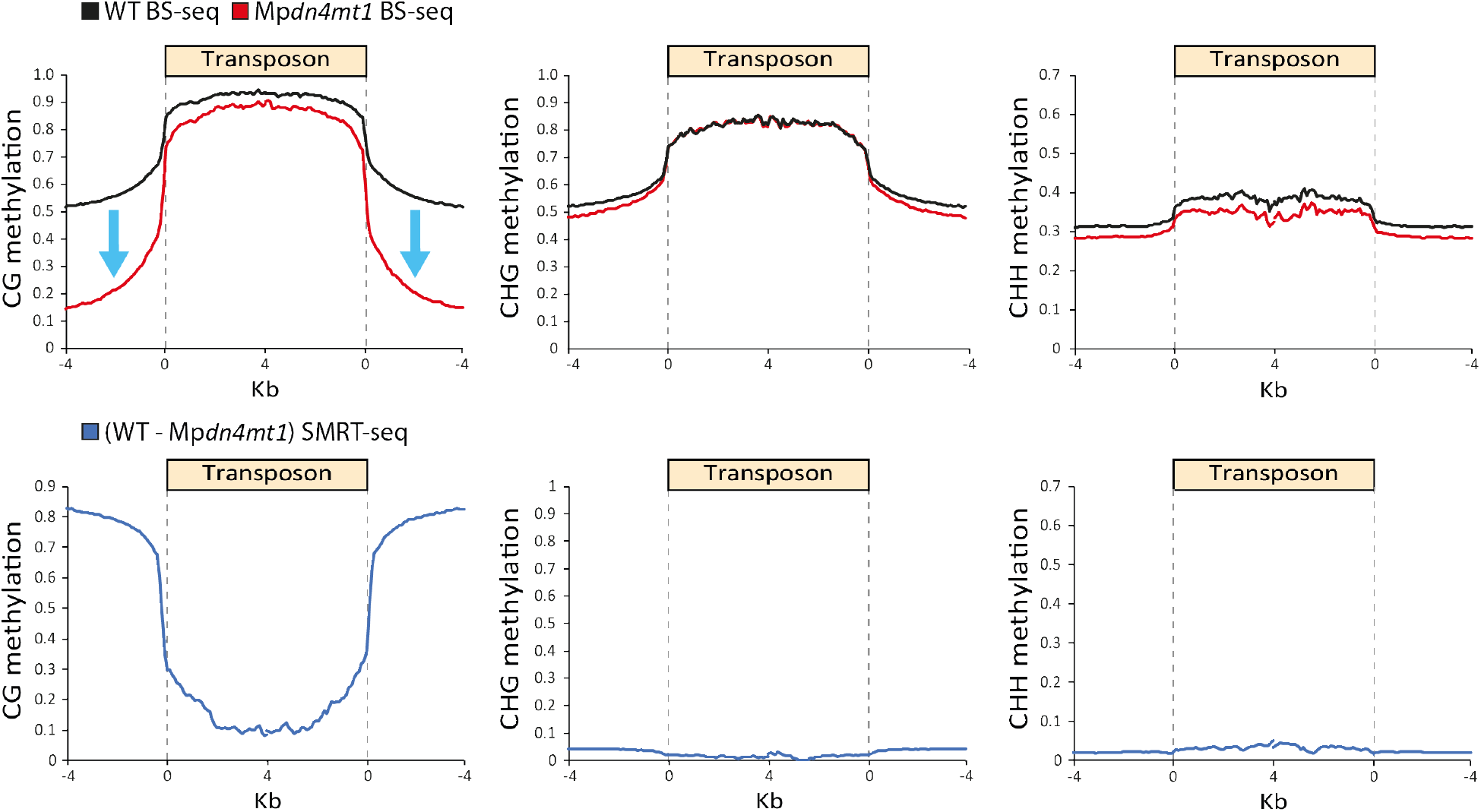
Cytosine methylation in WT and Mp*dn4mt1* mutant sperm measured by BS-seq and SMRT-seq. Transposons were aligned at the 5’ and 3’ ends (dashed lines) and average methylation levels for each 100-bp interval are plotted. The dashed line at zero represents the point of alignment. Blue arrows indicate loss of CG methylation outside of transposons in Mp*dn4mt1* mutant sperm.

**Fig. S9.**
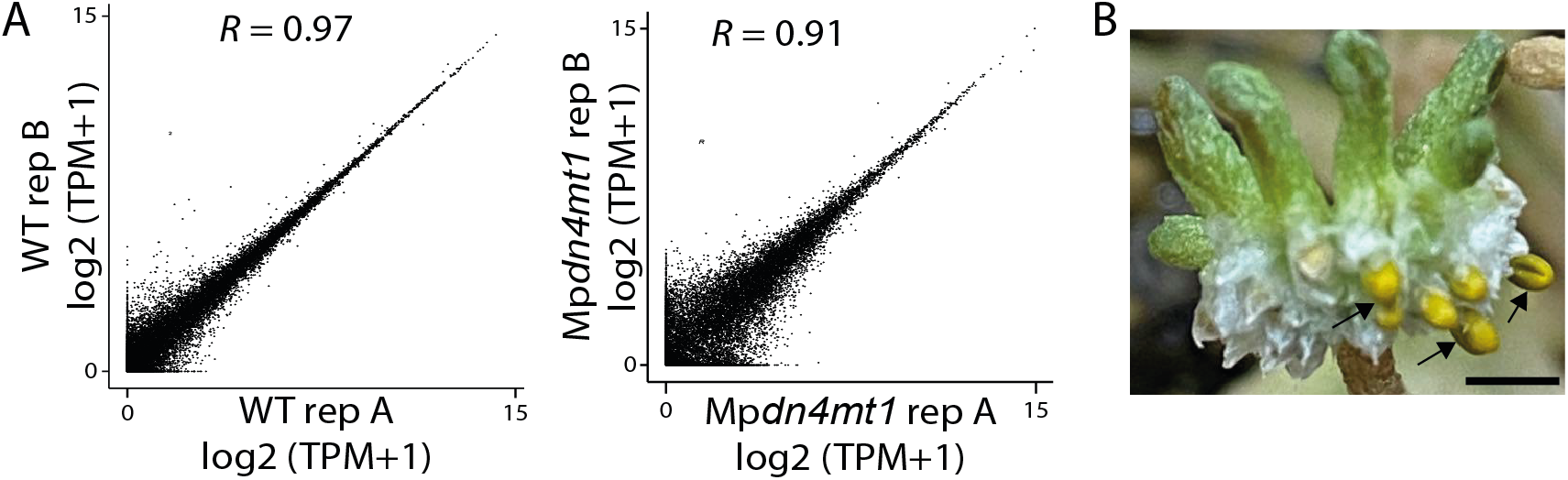
Functional and transcriptomic characterization of Mp*dn4mt1* mutant sperm. **(A)** Scatterplots showing high correlations between RNA-seq biological replicates (rep) of WT or Mp*dn4mt1* mutant sperm. *R*, Pearson’s coefficient. (**B**) Production of yellow sporangia (indicated by black arrows), which mark successful fertilization, on female WT plants after crossing with Mp*dn4mt1* sperm. Scale bar, 2.5 mm.

**Movie S1.** Swimming of wild-type sperm in water with example tracks marked. The movie was taken with an Axio Imager (Zeiss) using dark field settings at the rate of 10 frames per second (fps) for 10s.

**Movie S2.** Swimming of Mp*dn4mt1* mutant sperm in water with example tracks indicated. The movie was taken with an Axio Imager (Zeiss) using dark field settings at the rate of 10 frames per second (fps) for 10s.

**Movie S3.** Example of a WT sperm taken from Movie S1.

**Movie S4.** Example of a Mp*dn4mt1* mutant sperm taken from Movie S2.

## Materials and Methods

### Plant material and growth conditions

Male and female *Marchantia polymorpha, L. subsp. ruderalis*, accessions Takaragaike-1 (Tak-1, male) and Takaragaike-2 (Tak-2, female), were used. Plants were grown on plates containing ½ × strength Gamborg’s B5 medium with 1% agar (Sigma) under constant light at 21 °C, and Tak-1 thallus tissue was collected from 2-week old plants. Plants were transferred to Jiffy 7 peat compost pellets (Greens Hydroponics UK) after 2 weeks on plates and placed in a growth chamber at 21 °C with 70% humidity under constant light with far-red irradiation for induction of sexual reproduction, as described previously (*19, 34*). Spores were sterilized with Milton’s sterilizing solution (Milton Pharmaceutical UK) as described (*17*). Untransformed spores were grown to sexual maturity as above and used as wild-type (WT) in comparisons to Mp*dn4mt1* knockout mutants in methylation and phenotypic studies. Male and female plants were determined with sex chromosome-specific markers, rbm27 and rhf72 (*35*) (Table S5).

### Sperm, antheridia and embryo extraction

Sperm were isolated and developing antheridia were manually dissected as described previously (*19, 21*). Individual antheridia were transversely cut in half using fine needles (0.5mm × 25 mm). One half was immediately frozen in liquid nitrogen, while the other half was fixed in fixation buffer (4% paraformaldehyde, PBS, pH 7.5, 0.1% Triton X-100, 0.1% Tween-20). After a series of ethanol dehydration steps followed by resin embedding using a Technovit 7100 kit (Electron Microscopy Sciences), the fixed antheridium halves were cut into 5 μm sections with a microtome 2250 (Leica) and stained with Toluidine blue (*36*). DNA and RNA were extracted simultaneously from the frozen antheridium halves and individually from sperm using the Dynabeads® mRNA DIRECT^TM^ Micro Kit (Invitrogen). Embryos were dissected from fertilized gametophores 7 days following sperm application and the surrounding pseudoperianths were removed. Thallus, sperm and embryo DNA was also extracted as described (*37*).

### Plasmid construction and transformation

The Mp*DNMT3b* (Mp5g09290) knockdown construct was generated by synthesizing an artificial microRNA (amiRNA) construct (JW004; Table S5, ENSA), based on MpMIR160 (*38*) and designed as described (*39*) to target to an exon of Mp*DNMT3b*, and thereafter inserted into MpGWB103 (*40*) using the Gateway system (Invitrogen). For the Mp*CMTa* (Mp6g08650) knockdown construct, the amiRNA sequence was amplified and modified with primers JW490, JW495, and JW539-542 (Table S5) to target an exon of Mp*CMTa* and inserted into MpGWB103.

To generate *pro*Mp*DN4MT1a:*Mp*DN4MT1a-Citrine* and *pro*Mp*DN4MT1b:*Mp*DN4MT1b-Citrine* constructs, Tak-1 genomic DNA for each gene (Mp6g18330 and Mp6g18340) plus 2-kb upstream sequences were amplified using primers YL176, YL257, YL178, and YL258 (Table S5), and inserted into the destination vector pMpGWB207 (*40*) using the Gateway system (Invitrogen).

For CRISPR/Cas9 deletion of Mp*DN4MT1a* and Mp*DN4MT1b*, double nicking was carried out as previously described (*41, 42*) using pMpGE017 (*43*). Briefly, four different gRNAs (Table S5) were cloned into the BsaI site of pMpGE_En04 vector or into the multiplex vectors pBC-GE12, pBC-GE23 or pBC-GE34 (*41, 42, 44*). The four gRNAs cassettes were then cloned into pMpGE017 binary vector by LR reaction (Invitrogen).

Gemmae (Mp*dnmt3b* and Mp*cmta* knockdown mutants) or F1 spores (Mp*dn4mt1* CRISPR/Cas9 deletion mutants and *pro*Mp*DN4MT1a/b:*Mp*DN4MT1a/b-Citrine* lines) were transformed with the above constructs using agrobacteria transformation (strain GV6620) and positive transformants were selected on hygromycin (Mp*dnmt3b* and Mp*cmta* knockdown mutants, and Mp*dn4mt1* knockout mutant) or gentamycin (*pro*Mp*DN4MT1a/b:*Mp*DN4MT1a/b-Citrine* lines), as described *(45, 46)*. The knockdown lines were checked by qPCR using antheridiophores as previously described (*14*), with primers JW778 and JW779 for Mp*cmta* knockdown, primers JW448 and JW449 for Mp*dnmt3b* knockdown, and primers JW438 and JW439 (Table S5) to compare expression to Mp*EF1α* (Mp3g23400). Mp*dn4mt1* deletion was detected via PCR with primers YL1 and YL2 and sequenced with YL1 (Table S5). An 8807-bp deletion (Chr6: 20,611,704 - 20,620,510) including the tandem repeats of Mp*DN4MT1a* and *1b* (Chr6: 20,613,540 – 20,619,372) was obtained except for the inclusion of a 9-bp sequence (AGGTTTCAA; Chr6: 20,612,442 – 20,612,450) outside Mp*DN4MT1s*.

### Sequencing-library construction and analysis

4mC and 5mC spike-in controls were generated from pUC19 (NEB) PCR product synthesized with N4-methyl-dCTP (4mdCTP; Trilink) using primers JZ294/JZ295 (Table S5) and genomic lambda cl857 DNA (Promega) methylated with 5mC CG methyltransferase M. *SssI* (NEB), respectively, and added to DNA samples prior to library construction where appropriate for testing bisulfite conversion, as previously described *(31)*.

For antheridia (two biological replicates each for early, middle, and late stage used in Fig. 1 and Figs. S1 and S2), sperm (one biological replicate used in Figs. 1 and 2 and Figs. S1 to S3), Mp*DNMT3b* and Mp*CMTa* knockdown sperm (one biological replicate each used in Fig. 2 and Fig. S3), a single-cell-based method for BS-seq library preparation was carried out as previously described (*47*). For one biological replicate each of thallus, embryo, WT sperm and Mp*dn4mt1* sperm, single-end BS-seq libraries were also constructed with Ovation Ultralow Methyl-Seq Library Systems (Nugen, 0336) and EpiTect Bisulfite Kit (Qiagen, 59104) kits according to manufacturer’s instructions, with the incorporation of either one (WT sperm and Mp*dn4mt1* mutant sperm used in Fig. 3 and Figs. S7 and S8) or two rounds (WT sperm used in Fig. S7, and thallus and embryo used in Fig. 1 and Fig. S2) of bisulfite conversion. RNA-seq library preparation was either simultaneously carried out from the same input as the single-cell method bisulfite libraries (antheridia) or separately (WT sperm and Mp*dn4mt1* sperm; two replicates each) as described (*48*). All libraries were sequenced on NextSeq 500 (Illumina). RNA-seq data for thallus was obtained from published sources (*21*).

For BS-seq, low-quality reads and potential adaptor sequences were first trimmed using TrimGalore (v0.4.2) with default parameters. Biological replicates for antheridia were combined and DNA methylation analysis was performed based on Bismark (*49*), with ambiguous reads mapped randomly. Methylation data was mapped to the *Marchantia* chromosome assembly (v5.1) (*22*) as well as to the pUC19 vector sequence and lambda cl857 genome where appropriate. For all analyses regarding the *Marchantia* genome, only autosomes and the male sex chromosome (ChrV) were considered.

WT and Mp*dn4mt1* mutant sperm libraries for SMRT-seq were prepared using the SMRTbell Express Prep kit (v2.0; Pacific Biosciences, Menlo Park, CA, USA) following the low DNA input workflow as described (*50*). Libraries were sequenced under CCS running mode with one Sequel II SMRT Cell for each library via DNA Link (Seoul, Korea). SMRT libraries were mapped to the *Marchantia* (v5.1) genome using ppbm2 (v1.1.0). Base modifications were detected using ipdSummary (v2.0) with the following flags set (--identify 4mC --methylFraction).

Transposon meta-analysis and 5mC methylation analysis was performed as previously described (*51*). The proportion of the *Marchantia* genome considered to be methylated in thallus and during sperm development was taken as the percentage of 100-bp genomic windows (w100s) with fractional methylation > 0.2 for CG, > 0.1 for CHG, > 0.05 for CHH, > 0.1 for non-CG, or >0.1 for all cytosines (Cmethyl) out of the total number of sequenced w100s in the respective context. For comparisons between WT and knockdown sperm, a fractional methylation level > 0.3 in each of the contexts was used. w100s were considered to be associated with repeats if they overlapped or were within 500-bp of the *Marchantia* (v5.1) transposon annotation. For comparisons to LC-MS, methylation levels from BS-seq data were calculated using the average fractional methylation of single cytosines from Cmethyl. For SMRT-seq data, the proportion of N4 methylated cytosines for each w100 was taken as the number of 4mC cytosines detected in WT minus Mp*dn4mt1* over the total number of cytosines within the w100 sequence. Violin plots were generated using w100s with at least ten informative sequenced cytosines in the respective context for BS-seq data while all w100s were considered for SMRT-seq data. For both violin plots and calculations regarding the fractional 5mC and 4mC methylation of repeats, only w100s overlapping the transposon annotation with CG fractional methylation > 0.5 in thallus were considered.

For RNA-seq, low-quality reads and potential adaptor sequences were first trimmed using TrimGalore (v0.4.2) with default parameters. RNA-seq reads were processed with kallisto (v0.43.0) and statistical analysis was carried out with the sleuth package (v0.30.0) using the *Marchantia* (v5.1) gene annotation. Transcripts > 10 TPM in either WT or Mp*dn4mt1* were retained and those with a -log10 (*P*-value) < 0.1 between the two genotypes were discarded, leaving a total of 5,999 genes.

### Violin plots and heat maps

For all the violin plots in this study, the plot shows the distribution of the data and its probability density, with the box enclosing the middle 50% of the distribution (the horizontal line marks the median).

Heat maps (otherwise referred to as circus plots; www.circos.ca) were generated with fractional CG, CHG, and CHH methylation of 10-kb genomic windows.

### Sequence alignment and phylogenetic analysis

Orthologous protein sequences representing 5mC and β-class 4mC methyltransferases were obtained *(26, 28)*. Protein sequences up to motif III were taken from 5mC methyltransferases and relocated onto the C-terminus. All sequences were aligned alongside the MpDN4MT1s with MAFFT (v7) using default settings (*52*). A phylogenetic tree was constructed with the neighbor joining method using 100 bootstraps. A multiple sequence alignment of MpDN4MT1a, MpDN4MT1b, and prokaryotic 4mC methyltransferases was obtained and shaded with BoxShade (v3.2) using default parameters (http://sourceforge.net/projects/boxshade/). Motifs were labelled according to published data (*26*).

### Expression microscopy

Antheridiophores of p*ro*Mp*DN4MT1a/b:*Mp*DN4MT1a/b-Citrine* lines were manually cut longitudinally and observed under a fluorescence stereo microscope (Leica M205FA) and fluorescence was captured with a Leica DFC310FX color camera illuminated by LED 470nm-GFP. Antheridia with Citrine signal were manually taken out and observed under a confocal microscope (Leica SP8X) with a 40x/1.30 oil objective.

### Liquid chromatography–mass spectrometry

Thallus, WT sperm, and Mp*dn4mt1* mutant sperm DNA was digested into deoxyribonucleosides as previously described (*53*). The digested samples were centrifuged through 0.2 μm filters and 5 μl was analyzed by LC-MS/MS (Waters Aquity UPLC and XevoTQS MS). Separation was on a 100 × 2.1 mm 1.7 μ Kinetex C18 column (Phenomenex) using the following gradient of methanol (solvent B) versus 0.1% formic acid buffered to pH 8.3 with ammonium hydroxide (solvent A), run at 40 °C: 0 min, 0% B; 2 min, 0% B; 4 min, 10% B; 4.5 min, 90% B; 5.5 min, 90% B; 5.6 min, 0% B; 9.6 min, 0% B. The flow rate was 0.5 mL·min^−1^ except during the periods 4.0 – 4.5 min and 5.5 – 6.5 min to avoid excessive pressure. Analytes were quantified against external standard curves by multiple reaction monitoring (MRM) detecting the following transitions from positive-mode electrospray: 228→112 (unmethylated C), 242→126 (4mC and 5mC), and 256→140 (4,5mC). 4mC and 5mC eluted at different retention times. Collision energies and cone voltages were optimized using Waters’ Intellistart software. Spray chamber conditions were 3.5 kV spray voltage, 500 °C desolvation temperature, 900 L·hr^−1^ desolvation gas, 150 L·hr^−1^ cone gas, and 7.0 bar nebulizer pressure. Deoxycytidine (Sigma), 5-Me-2’-deoxycytidine (Cayman chemicals), 2’-Deoxy-N4-methylcytidine (CarboSynth), and N4,5-Dimethyldeoxycytidine (Toronto Research Chemicals) were used as standards.

### Phenotyping of Mp*dn4mt1* mutant sperm

Swimming dynamics of *Marchantia* sperm were examined as previously described (*54*), with modifications. WT and Mp*dn4mt1* mutant sperm discharged in water were observed under an Axio Imager Z2 (Zeiss) with Hamamatsu camera and 40x/0.75 air objective in dark field and video-recorded at the rate of 10 frames per second (fps) for 10s. To track sperm, TrackMate (v6.0.1) (*55*), a plugin bundled in ImageJ (*56*) (v1.53c; the Fiji distribution) was used, *n* = 30 (WT), *n* = 28 (Mp*dn4mt1*). Swimming velocity was calculated by track displacement divided by track duration time. Directionality was calculated by track displacement divided by sum of links displacement.

## References

1. R. J. Schmitz, Z. A. Lewis, M. G. Goll, DNA methylation: shared and divergent features across eukaryotes. Trends Genet. 35, 818–827 (2019).

2. M. A. Sánchez-Romero, J. Casadesús, The bacterial epigenome. Nat. Rev. Microbiol. 18, 7–20 (2020).

3. J. Beaulaurier, E. E. Schadt, G. Fang, Deciphering bacterial epigenomes using modern sequencing technologies. Nat. Rev. Genet. 20, 157–172 (2019).

4. P. Ye, Y. Luan, K. Chen, Y. Liu, C. Xiao, Z. Xie, MethSMRT: an integrative database for DNA N6-methyladenine and N4-methylcytosine generated by single-molecular real-time sequencing. Nucleic Acids Res. 45, D85’D89 (2017).

5. Z. K. O’Brown, K. Boulias, J. Wang, S. Y. Wang, N. M. O’Brown, Z. Hao, H. Shibuya, P.-E. Fady, Y. Shi, C. He, S. G. Megason, T. Liu, E. L. Greer, Sources of artifact in measurements of 6mA and 4mC abundance in eukaryotic genomic DNA. BMC Genom. 20, 445 (2019).

6. A. de Mendoza, R. Lister, O. Bogdanovic, Evolution of DNA methylome diversity in eukaryotes. J. Mol. Biol. 432, 1687–1705 (2020).

7. A. Zemach, I. E. McDaniel, P. Silva, D. Zilberman, Genome-wide evolutionary analysis of eukaryotic DNA methylation. Science 328, 916–919 (2010).

8. J. A. Law, S. E. Jacobsen, Establishing, maintaining and modifying DNA methylation patterns in plants and animals. Nat. Rev. Genet. 11, 204–220 (2010).

9. C. Luo, P. Hajkova, J. R. Ecker, Dynamic DNA methylation: In the right place at the right time. Science 361, 1336–1340 (2018).

10. H. Zhang, Z. Lang, J.-K. Zhu, Dynamics and function of DNA methylation in plants. Nat. Rev. Mol. Cell Biol. 19, 489–506 (2018).

11. C. E. Niederhuth, A. J. Bewick, L. Ji, M. S. Alabady, K. D. Kim, Q. Li, N. A. Rohr, A. Rambani, J. M. Burke, J. A. Udall, C. Egesi, J. Schmutz, J. Grimwood, S. A. Jackson, N. M. Springer, R. J. Schmitz, Widespread natural variation of DNA methylation within angiosperms. Genome Biol. 17, 194 (2016).

12. A. P. Feinberg, M. A. Koldobskiy, A. Göndör, Epigenetic modulators, modifiers and mediators in cancer aetiology and progression. Nat. Rev. Genet. 17, 284–299 (2016).

13. M. V. C. Greenberg, D. Bourc’his, The diverse roles of DNA methylation in mammalian development and disease. Nat. Rev. Mol. Cell Biol. 20, 590–607 (2019).

14. J. Walker, H. Gao, J. Zhang, B. Aldridge, M. Vickers, J. D. Higgins, X. Feng, Sexual-lineage-specific DNA methylation regulates meiosis in *Arabidopsis*. Nat. Genet. 50, 130–137 (2018).

15. S. Seisenberger, J. R. Peat, T. A. Hore, F. Santos, W. Dean, W. Reik, Reprogramming DNA methylation in the mammalian life cycle: building and breaking epigenetic barriers. Philos. Trans. R. Soc. Lond., B, Biol. Sci. 368, 20110330 (2013).

16. W. W. Tang, T. Kobayashi, N. Irie, S. Dietmann, M. A. Surani, Specification and epigenetic programming of the human germ line. Nat. Rev. Genet. 17, 585–600 (2016).

17. J. L. Bowman, T. Kohchi, K. T. Yamato, J. Jenkins, S. Shu, K. Ishizaki, S. Yamaoka, R. Nishihama, Y. Nakamura, F. Berger, C. Adam, S. S. Aki, F. Althoff, T. Araki, M. A. Arteaga-Vazquez, S. Balasubrmanian, K. Barry, D. Bauer, C. R. Boehm, L. Briginshaw, J. Caballero-Perez, B. Catarino, F. Chen, S. Chiyoda, M. Chovatia, K. M. Davies, M. Delmans, T. Demura, T. Dierschke, L. Dolan, A. E. Dorantes-Acosta, D. M. Eklund, S. N. Florent, E. Flores-Sandoval, A. Fujiyama, H. Fukuzawa, B. Galik, D. Grimanelli, J. Grimwood, U. Grossniklaus, T. Hamada, J. Haseloff, A. J. Hetherington, A. Higo, Y. Hirakawa, H. N. Hundley, Y. Ikeda, K. Inoue, S.-i. Inoue, S. Ishida, Q. Jia, M. Kakita, T. Kanazawa, Y. Kawai, T. Kawashima, M. Kennedy, K. Kinose, T. Kinoshita, Y. Kohara, E. Koide, K. Komatsu, S. Kopischke, M. Kubo, J. Kyozuka, U. Lagercrantz, S.-S. Lin, E. Lindquist, A. M. Lipzen, C.-W. Lu, E. De Luna, R. A. Martienssen, N. Minamino, M. Mizutani, M. Mizutani, N. Mochizuki, I. Monte, R. Mosher, H. Nagasaki, H. Nakagami, S. Naramoto, K. Nishitani, M. Ohtani, T. Okamoto, M. Okumura, J. Phillips, B. Pollak, A. Reinders, M. Rövekamp, R. Sano, S. Sawa, M. W. Schmid, M. Shirakawa, R. Solano, A. Spunde, N. Suetsugu, S. Sugano, A. Sugiyama, R. Sun, Y. Suzuki, M. Takenaka, D. Takezawa, H. Tomogane, M. Tsuzuki, T. Ueda, M. Umeda, J. M. Ward, Y. Watanabe, K. Yazaki, R. Yokoyama, Y. Yoshitake, I. Yotsui, S. Zachgo, J. Schmutz, Insights into land plant evolution garnered from the *Marchantia polymorpha* genome. Cell 171, 287–304.e215 (2017).

18. S. Takuno, J.-H. Ran, B. S. Gaut, Evolutionary patterns of genic DNA methylation vary across land plants. Nat. Plants 2, 15222 (2016).

19. M. W. Schmid, A. Giraldo-Fonseca, M. Rövekamp, D. Smetanin, J. L. Bowman, U. Grossniklaus, Extensive epigenetic reprogramming during the life cycle of *Marchantia polymorpha*. Genome Biol. 19, 9 (2018).

20. A. J. Bewick, C. E. Niederhuth, L. Ji, N. A. Rohr, P. T. Griffin, J. Leebens-Mack, R. J. Schmitz, The evolution of CHROMOMETHYLASES and gene body DNA methylation in plants. Genome Biol. 18, 65 (2017).

21. A. Higo, M. Niwa, K. T. Yamato, L. Yamada, H. Sawada, T. Sakamoto, T. Kurata, M. Shirakawa, M. Endo, S. Shigenobu, K. Yamaguchi, K. Ishizaki, R. Nishihama, T. Kohchi, T. Araki, Transcriptional framework of male gametogenesis in the liverwort *Marchantia polymorpha* L. Plant Cell Physiol. 57, 325–338 (2016).

22. S. A. Montgomery, Y. Tanizawa, B. Galik, N. Wang, T. Ito, T. Mochizuki, S. Akimcheva, J. L. Bowman, V. Cognat, L. Maréchal-Drouard, H. Ekker, S.-F. Hong, T. Kohchi, S.-S. Lin, L.-Y. D. Liu, Y. Nakamura, L. R. Valeeva, E. V. Shakirov, D. E. Shippen, W.-L. Wei, M. Yagura, S. Yamaoka, K. T. Yamato, C. Liu, F. Berger, Chromatin organization in early land plants reveals an ancestral association between H3K27me3, transposons, and constitutive heterochromatin. Curr. Biol. 30, 573–588.e577 (2020).

23. R. Yaari, A. Katz, K. Domb, K. D. Harris, A. Zemach, N. Ohad, RdDM-independent de novo and heterochromatin DNA methylation by plant CMT and DNMT3 orthologs. Nat. Commun. 10, 1613 (2019).

24. C. Noy-Malka, R. Yaari, R. Itzhaki, A. Mosquna, N. Auerbach Gershovitz, A. Katz, N. Ohad, A single CMT methyltransferase homolog is involved in CHG DNA methylation and development of *Physcomitrella patens*. Plant Mol. Biol. 84, 719–735 (2014).

25. Y. Ikeda, R. Nishihama, S. Yamaoka, M. A. Arteaga-Vazquez, A. Aguilar-Cruz, D. Grimanelli, R. Pogorelcnik, R. A. Martienssen, K. T. Yamato, T. Kohchi, T. Hirayama, O. Mathieu, Loss of CG methylation in *Marchantia polymorpha* causes disorganization of cell division and reveals unique DNA methylation regulatory mechanisms of non-CG methylation. Plant Cell Physiol. 59, 2421–2431 (2018).

26. J. M. Bujnicki, M. Radlinska, Molecular evolution of DNA-(cytosine-N4) methyltransferases: evidence for their polyphyletic origin. Nucleic Acids Res. 27, 4501–4509 (1999).

27. S. Klimasauskas, A. Timinskas, S. Menkevicius, D. Butkiene, V. Butkus, A. Janulaitis, Sequence motifs characteristic of DNA[cytosine-N4]methyltransferases: similarity to adenine and cytosine-C5 DNA-methylases. Nucleic Acids Res. 17, 9823–9832 (1989).

28. T. P. Jurkowski, A. Jeltsch, On the evolutionary origin of eukaryotic DNA methyltransferases and Dnmt2. PLoS One 6, e28104 (2011).

29. G. Vilkaitis, S. Klimasauskas, Bisulfite sequencing protocol displays both 5-methylcytosine and N4-methylcytosine. Anal. Biochem. 271, 116–119 (1999).

30. M. Frommer, L. E. McDonald, D. S. Millar, C. M. Collis, F. Watt, G. W. Grigg, P. L. Molloy, C. L. Paul, A genomic sequencing protocol that yields a positive display of 5-methylcytosine residues in individual DNA strands. Proc. Natl. Acad. Sci. U.S.A. 89, 1827–1831 (1992).

31. M. Yu, L. Ji, D. A. Neumann, D. H. Chung, J. Groom, J. Westpheling, C. He, R. J. Schmitz, Base-resolution detection of N4-methylcytosine in genomic DNA using 4mC-Tet-assisted-bisulfite-sequencing. Nucleic Acids Res. 43, e148 (2015).

32. B. M. Davis, M. C. Chao, M. K. Waldor, Entering the era of bacterial epigenomics with single molecule real time DNA sequencing. Curr. Opin. Microbiol. 16, 192–198 (2013).

33. T. A. Clark, I. A. Murray, R. D. Morgan, A. O. Kislyuk, K. E. Spittle, M. Boitano, A. Fomenkov, R. J. Roberts, J. Korlach, Characterization of DNA methyltransferase specificities using single-molecule, real-time DNA sequencing. Nucleic Acids Res. 40, e29 (2012).

34. S. Chiyoda, K. Ishizaki, H. Kataoka, K. T. Yamato, T. Kohchi, Direct transformation of the liverwort *Marchantia polymorpha* L. by particle bombardment using immature thalli developing from spores. Plant Cell Rep. 27, 1467 (2008).

35. M. Fujisawa, K. Hayashi, T. Nishio, T. Bando, S. Okada, K. T. Yamato, H. Fukuzawa, K. Ohyama, Isolation of X and Y chromosome-specific DNA markers from a liverwort, *Marchantia polymorpha*, by representational difference analysis. Genetics 159, 981–985 (2001).

36. X. Feng, H. G. Dickinson, Tapetal cell fate, lineage and proliferation in the *Arabidopsis* anther. Development 137, 2409–2416 (2010).

37. A. Healey, A. Furtado, T. Cooper, R. J. Henry, Protocol: a simple method for extracting next-generation sequencing quality genomic DNA from recalcitrant plant species. Plant Methods 10, 21 (2014).

38. E. Flores-Sandoval, T. Dierschke, T. J. Fisher, J. L. Bowman, Efficient and inducible use of artificial microRNAs in *Marchantia polymorph*a. Plant Cell Physiol. 57, 281–290 (2016).

39. V. A. Jones, L. Dolan, MpWIP regulates air pore complex development in the liverwort *Marchantia polymorpha*. Development 144, 1472–1476 (2017).

40. K. Ishizaki, R. Nishihama, M. Ueda, K. Inoue, S. Ishida, Y. Nishimura, T. Shikanai, T. Kohchi, Development of gateway binary vector series with four different selection markers for the liverwort *Marchantia polymorpha*. PLoS One 10, e0138876 (2015).

41. F. A. Ran, P. D. Hsu, C. Y. Lin, J. S. Gootenberg, S. Konermann, A. E. Trevino, D. A. Scott, A. Inoue, S. Matoba, Y. Zhang, F. Zhang, Double nicking by RNA-guided CRISPR Cas9 for enhanced genome editing specificity. Cell 154, 1380–1389 (2013).

42. B. Shen, W. Zhang, J. Zhang, J. Zhou, J. Wang, L. Chen, L. Wang, A. Hodgkins, V. Iyer, X. Huang, W. C. Skarnes, Efficient genome modification by CRISPR-Cas9 nickase with minimal off-target effects. Nat. Methods 11, 399–402 (2014).

43. I. Monte, S. Ishida, A. M. Zamarreño, M. Hamberg, J. M. Franco-Zorrilla, G. García-Casado, C. Gouhier-Darimont, P. Reymond, K. Takahashi, J. M. García-Mina, R. Nishihama, T. Kohchi, R. Solano, Ligand-receptor co-evolution shaped the jasmonate pathway in land plants. Nat. Chem. Biol. 14, 480–488 (2018).

44. T. Sakuma, A. Nishikawa, S. Kume, K. Chayama, T. Yamamoto, Multiplex genome engineering in human cells using all-in-one CRISPR/Cas9 vector system. Sci. Rep. 4, 5400 (2014).

45. S. Tsuboyama, Y. Kodama, AgarTrap: a simplified Agrobacterium-mediated transformation method for sporelings of the liverwort *Marchantia polymorpha* L. Plant Cell Physiol. 55, 229–236 (2014).

46. S. Tsuboyama, S. Nonaka, H. Ezura, Y. Kodama, Improved G-AgarTrap: A highly efficient transformation method for intact gemmalings of the liverwort *Marchantia polymorpha*. Sci. Rep. 8, 10800 (2018).

47. S. A. Smallwood, H. J. Lee, C. Angermueller, F. Krueger, H. Saadeh, J. Peat, S. R. Andrews, O. Stegle, W. Reik, G. Kelsey, Single-cell genome-wide bisulfite sequencing for assessing epigenetic heterogeneity. Nat. Methods 11, 817–820 (2014).

48. S. Picelli, O. R. Faridani, Å. K. Björklund, G. Winberg, S. Sagasser, R. Sandberg, Full-length RNA-seq from single cells using Smart-seq2. Nat. Protoc. 9, 171–181 (2014).

49. F. Krueger, S. R. Andrews, Bismark: a flexible aligner and methylation caller for Bisulfite-Seq applications. Bioinformatics 27, 1571–1572 (2011).

50. S. B. Kingan, H. Heaton, J. Cudini, C. C. Lambert, P. Baybayan, B. D. Galvin, R. Durbin, J. Korlach, M. K. N. Lawniczak, A high-quality de novo genome assembly from a single mosquito using PacBio sequencing. Genes (Basel) 10, (2019).

51. C. A. Ibarra, X. Feng, V. K. Schoft, T.-F. Hsieh, R. Uzawa, J. A. Rodrigues, A. Zemach, N. Chumak, A. Machlicova, T. Nishimura, D. Rojas, R. L. Fischer, H. Tamaru, D. Zilberman, Active DNA demethylation in plant companion cells reinforces transposon methylation in gametes. Science 337, 1360–1364 (2012).

52. K. Katoh, J. Rozewicki, K. D. Yamada, MAFFT online service: multiple sequence alignment, interactive sequence choice and visualization. Brief. Bioinform. 20, 1160–1166 (2017).

53. E. P. Quinlivan, J. F. Gregory, DNA digestion to deoxyribonucleoside: A simplified one-step procedure. Anal. Biochem. 373, 383–385 (2008).

54. A. Higo, T. Kawashima, M. Borg, M. Zhao, I. López-Vidriero, H. Sakayama, S. A. Montgomery, H. Sekimoto, D. Hackenberg, M. Shimamura, T. Nishiyama, K. Sakakibara, Y. Tomita, T. Togawa, K. Kunimoto, A. Osakabe, Y. Suzuki, K. T. Yamato, K. Ishizaki, R. Nishihama, T. Kohchi, J. M. Franco-Zorrilla, D. Twell, F. Berger, T. Araki, Transcription factor DUO1 generated by neo-functionalization is associated with evolution of sperm differentiation in plants. Nat. Commun. 9, 5283 (2018).

55. J.-Y. Tinevez, N. Perry, J. Schindelin, G. M. Hoopes, G. D. Reynolds, E. Laplantine, S. Y. Bednarek, S. L. Shorte, K. W. Eliceiri, TrackMate: An open and extensible platform for single-particle tracking. Methods 115, 80–90 (2017).

56. J. Schindelin, I. Arganda-Carreras, E. Frise, V. Kaynig, M. Longair, T. Pietzsch, S. Preibisch, C. Rueden, S. Saalfeld, B. Schmid, J.-Y. Tinevez, D. J. White, V. Hartenstein, K. Eliceiri, P. Tomancak, A. Cardona, Fiji: an open-source platform for biological-image analysis. Nat. Methods 9, 676–682 (2012).

